# NeuroCDS: Integrating Local and Global Neural Network Representations via Structural Constrained Viterbi Decoding for Robust CDS Annotation

**DOI:** 10.64898/2026.04.29.721598

**Authors:** Ziheng Mei, Zhexi Xie, Lingrui Wu, Chao Wei

## Abstract

**Motivation:** Robust annotation of Coding Sequences (CDS) is critical for downstream transcriptomics, yet heavily fragmented *de novo* RNA-Seq assemblies pose a severe challenge. Traditional computational tools rely on fixed, hand-crafted features that are prone to fail when canonical sequence signals are truncated. While recent deep learning models excel at automatically extracting complex representations, they predominantly treat these as isolated prediction tasks. Lacking a joint inference mechanism to enforce structural constraints, existing models occasionally output biologically invalid predictions. Therefore, a computational framework capable of fusing heterogeneous neural network representations for joint annotation is critically needed.

**Results:** We present NeuroCDS, a reliable framework that bridges the effective representation capabilities of deep neural networks with the structural rigor of dynamic programming. NeuroCDS employs a dual-branch architecture: a Convolutional Neural Network (CNN) acts as a local sensor to extract Translation Initiation Sites (TIS), while a Temporal Convolutional Network (TCN) acts as a global sensor to evaluate continuous regional coding potential. The primary contribution of NeuroCDS lies in a structurally constrained Viterbi Decoding algorithm designed to fuse these heterogeneous signals. This joint inference mechanism strictly enforces biological grammars (e.g., reading frame preservation) to dynamically calculate the globally optimal transcript structure via a tripartite state space. Crucially, by introducing a dynamic length normalization mechanism, our formulation adaptively leverages global continuous representations to stably annotate both intact transcripts and highly truncated fragments. Comprehensive evaluations demonstrate that NeuroCDS achieves high F1-scores on full-length transcripts and maintains robust performance on complex Ribo-seq validated datasets, comparing favorably against traditional HMM-based and heuristic methodologies.

**Availability:** Source code, pre-trained models, and datasets are freely available at https://github.com/hgcwei/NeuroCDS.

## Introduction

High-throughput RNA sequencing (RNA-Seq) has revolutionized our ability to profile the transcribed genome across diverse biological conditions [1, 2]. For organisms lacking a high-quality reference genome, *de novo* transcriptome assembly has become the indispensable and most cost-effective approach for investigating gene expression and discovery [3, 4]. However, a critical downstream bottleneck in analyzing these newly reconstructed transcriptomes is the stable annotation of coding sequences (CDS), which serves as the prerequisite for translating nucleotide sequences into functional protein products. Despite continuous algorithmic advancements in assembly strategies [4–6], *de novo* transcriptomes frequently suffer from technical artifacts. Sequencing errors, uneven read coverage, and complex alternative splicing isoforms often lead to highly fragmented contigs [7, 8]. A substantial proportion of these generated contigs represent only partial transcripts truncated at the 5’ end, the 3’ end, or both. This pervasive fragmentation poses a severe challenge for reliably identifying functional protein-coding regions.

To address this challenge, various computational efforts have been proposed over the years. Traditional computational tools have predominantly relied on manually engineered sequence features and classical statistical frameworks to delineate open reading frames (ORFs). For instance, widely utilized tools depend on distinct heuristic and statistical signatures: TransDecoder [5] prioritizes the longest ORF and typically employs Position-Specific Scoring Matrices (PSSMs) to evaluate localized sequence motifs, such as translation initiation and termination signatures; CPAT [9] utilizes a logistic regression framework to integrate multiple hand-crafted sequence metrics, including hexamer usage bias, the Fickett TESTCODE score, and ORF length; while GeneMarkS-T [10] builds species-specific inhomogeneous Markov models based on *k*-mer frequencies to represent coding and non-coding sequences. Furthermore, to specifically address sequence fragmentation and sequencing artifacts, tools like ESTScan [11] and CodAn [12] employ profile Hidden Markov Models (HMMs) and Generalized HMMs (GHMMs) [13]. These models utilize pre-calculated transition penalties and static emission tables—such as codon or hexamer probabilities—to parse sequences into discrete states (e.g., 5’ UTR, CDS, 3’ UTR). Despite their historical utility, these conventional approaches exhibit inherent algorithmic bottlenecks. Primarily, their reliance on explicit, predefined feature sets severely limits their capacity for the deep mining of highly complex, non-linear sequence patterns. They struggle to adequately enforce label consistency across extended coding bodies and fail to capture intricate, long-range nucleotide dependencies beyond their fixed *k*-mer windows. Consequently, their predictive performance exhibits substantial fluctuations when applied to heavily fragmented contigs where canonical boundary signals (such as a clear Kozak sequence or intact start/stop codons) are truncated or obscured.

In recent years, deep learning methodologies have been introduced to overcome the limitations of manual feature engineering, driving significant advancements in bioinformatics [14, 15]. By deploying diverse neural architectures, researchers have successfully isolated and modeled distinct biological sequence signals. For instance, Convolutional Neural Networks (CNNs) have proven highly effective at capturing localized spatial motifs; tools such as TISRover [16] leverage deep CNNs to precisely identify Translation Initiation Sites (TIS) by automatically learning the complex non-linear patterns (e.g., Kozak consensus) surrounding the start codon. Conversely, capturing the continuous properties of Open Reading Frames (ORFs) requires architectures with expansive receptive fields. Recent advancements, such as NeuroTIS+ [17], have utilized Temporal Convolutional Networks (TCNs) integrated with codon usage frequencies to evaluate regional coding potential, effectively incorporating label consistency constraints to smooth predictions across the reading frame. However, despite their superiority in extracting specific, non-linear sequence features, current neural network approaches possess notable architectural drawbacks when applied to full-scale transcriptomic annotation. Most fundamentally, they predominantly treat the identification of local motifs (e.g., TIS) and the evaluation of regional coding bodies as isolated, independent prediction tasks, lacking a unified global perspective. Without a mathematically rigorous joint inference mechanism to enforce structural constraints and biological grammars, purely neural models occasionally output biologically invalid predictions (e.g., overlapping reading frames or unhandled in-frame stop codons). Furthermore, existing deep learning models generally lack specialized modeling strategies tailored for sequence fragmentation, rendering their predictions unstable when applied to the heavily truncated assemblies typical of real-world *de novo* transcriptomics.

Recognizing the respective limitations of both traditional statistical frameworks (weak feature representation) and current deep learning methods (lack of global structural consensus and fragment adaptability), we present NeuroCDS, a reliable computational framework designed to synergize the powerful representation capabilities of deep neural networks with the structural rigor of dynamic programming. Building upon the foundational logic of both algorithmic paradigms, our framework conceptualizes CDS prediction as a holistic, structurally constrained sequence parsing problem. It employs a dual-branch neural architecture: a CNN acts as a dependable “local sensor” to extract Translation Initiation Sites (TIS), while a TCN acts as a continuous “global sensor” to evaluate regional coding potential. These heterogeneous neural representations are then systematically integrated via a custom dynamic programming algorithm, effectively bridging deep learning representations with mathematically advantageous structural joint annotation.

The primary contributions of this paper are summarized as follows:

- **Deep Mining of Sequence Features:** We propose a synergistic dual-branch neural architecture (CNN and TCN) that automatically mines complex sequence features, effectively capturing both localized spatial motifs and long-range nucleotide dependencies without manual feature engineering.
- **Integration of Global Structural Information:** We introduce a structurally constrained Segmental Viterbi Decoding algorithm that transforms isolated neural predictions into a globally optimal transcript structure. This joint inference mechanism strictly enforces biological grammars, such as reading frame consistency, to eliminate biologically invalid outputs.
- **Specialized Modeling for Sequence Fragments:** We design a dynamic length normalization mechanism to specifically address the problem of sequence fragmentation. This allows the framework to adaptively leverage global continuous representations to stably annotate both intact transcripts and highly truncated fragments (e.g., sequences missing canonical start or stop codons).
- **Comprehensive Robustness and Generalization:** Extensive evaluations demonstrate that NeuroCDS achieves state-of-the-art performance on full-length transcripts and complex Ribo-seq validated datasets. It successfully suppresses false positives in non-coding artifacts (e.g., 3’ UTRs and ncRNAs) and exhibits strong cross-species generalization potential.

## Materials and Methods

### Datasets and Problem Formulation

To ensure a rigorous and fair comparison with state-of-the-art methods, we trained and evaluated NeuroCDS using the comprehensive benchmark datasets originally established by the *CodAn* study [12]. Specifically, our evaluation employed the entirety of the datasets provided by CodAn, which encompass representative sequences across major eukaryotic clades, including Vertebrates, Invertebrates, Plants, and Fungi. Furthermore, to comprehensively evaluate the model’s robustness against sequence length degradation and its ability to suppress non-coding noise, we expanded the experimental data by additionally incorporating high-quality, full-length transcripts and non-coding sequences for human (*Homo sapiens*) and mouse (*Mus musculus*) downloaded directly from the NCBI RefSeq database.

All constructed datasets were strictly partitioned into independent training and testing sets at the species level to entirely prevent data leakage. The training sets comprise intact, full-length representative transcripts. Conversely, comprehensive testing sets incorporate both full-length transcripts and systematically generated 5’-partial, 3’-partial, and internal transcript fragments, allowing us to rigorously assess the framework’s robustness against typical *de novo* assembly artifacts.

Mathematically, we formulate the CDS annotation task as a structural sequence parsing problem. Let *S* = (*s*_1_, *s*_2_, …, *s*_*L*_) denote an input RNA sequence of length *L*. The objective is to extract a valid coding interval denoted by indices (*i, j*), where 1 ≤*i < j* ≤*L*, corresponding to the valid translation initiation and termination sites. A valid segment must strictly satisfy two biological grammars: its length must be a multiple of three (formulated as (*j* − *i* + 1) (mod 3) = 0), and the internal reading frame must not contain any in-frame stop codons. Our goal is to deduce the optimal structural segment 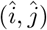by maximizing the joint probability of local TIS representations and continuous global coding potential.

### Neural Emission Probability Estimators

Given the established efficacy of deep learning architectures in genomic feature extraction, NeuroCDS employs standard convolutional models to compute preliminary emission probabilities, which serve as foundational inputs for our downstream decoding framework.

### Local TIS Representation via CNN

To capture the highly localized spatial motifs characterizing translation initiation signatures (e.g., the Kozak consensus sequence), we utilize a deep 1D Convolutional Neural Network (CNN). By sliding a defined sequence context window across the one-hot encoded transcript, the CNN autonomously extracts non-linear local dependencies. Following hierarchical feature extraction through max-pooling and dense layers, two parallel CNN modules independently output continuous position-wise probability arrays: *P*_*start*_ ∈ [0, 1]^*L*^. These arrays act as the discrete local emission representations, identifying putative TIS prediction spikes.

### Global Regional Coding Potential Representation via TCN

While the CNN is effective at localizing TIS, evaluating the continuous coding body of an open reading frame requires an expansive receptive field. For this, we utilize a Temporal Convolutional Network (TCN). To enforce biological relevance, the TCN processes an explicitly constructed overlapping *Codon Matrix* rather than single nucleotides, directly capturing synonymous codon usage biases. The input sequences are parsed using the Biopython package, converting raw nucleotides into a 64-dimensional feature vector representing the frequency of all possible codons.

Specifically, the TCN architecture in NeuroCDS consists of two residual stacks (nb_stacks = 2). Each stack contains a series of 1D convolutional layers with 15 filters (nb_filters = 15) and a large kernel size of 20 (kernel_size = 20). To efficiently capture long-range dependencies across the transcript without a massive increase in parameters, we implement dilated convolutions with a progressive expansion pattern of [1, 2, 4]. This configuration ensures that the global sensor can evaluate coding potential across expansive regional contexts.

A persistent challenge in CDS prediction is the generation of high-frequency positional noise. Because a valid CDS is translated continuously in units of three, we explicitly incorporate Consistency Modeling. For a specific coordinate *t*, the smoothed global coding potential *P*_*coding*_(*t*) is constrained by computing the average logit of its intra-codon neighbors within the corresponding reading frame:

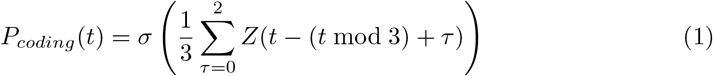

where *Z* represents the raw TCN output logits and *σ* is the sigmoid function. This biologically motivated constraint effectively eradicates intra-codon noise, forcing the TCN to output a structurally smooth, continuous signal that effectively complements the discrete TIS prediction spikes generated by the CNN.

### Structural Parsing via Structurally Constrained Viterbi Decoding

The primary algorithmic innovation of NeuroCDS lies in formulating the parsing of structural predictions not as independent thresholding events, but as a mathematically reliable sequence decoding problem. Rather than evaluating individual nucleotides, our algorithm processes the sequence strictly in units of triplets (codons) for each of the three potential reading frames independently, natively ensuring the structural constraint of reading frame preservation.

### Tripartite State Space and Biological Grammar Constraints

For each reading frame, we define a comprehensive tripartite state space: State 0 (*S*_0_) represents the 5’ non-coding region (5’ UTR), State 1 (*S*_1_) defines the active coding sequence (CDS), and State 2 (*S*_2_) denotes the 3’ non-coding region (3’ UTR).

Transitions between these discrete states are strictly governed by biological grammar. The transition from *S*_0_ to *S*_1_ requires a putative Translation Initiation Site (TIS) event, whereas transitions out of *S*_1_ correspond to translation termination. Crucially, the presence of an in-frame stop codon (TAA, TAG, TGA) acts as an absolute physical roadblock. Furthermore, to minimize the impact of random short ORFs on the prediction, we implemented a structural length constraint (e.g., a minimum of 90 base pairs).

### Dynamic Programming Formulation and Algorithmic Complexity

To systematically fuse the discrete localized CNN signals and the continuous TCN coding potential, we deploy a Viterbi dynamic programming algorithm. Let *N* be the total number of valid codons in a given reading frame. We receive three streams of neural emission logits evaluated at each codon index *t* ∈ {1, 2, …, *N*}: the positive coding potential ℰ_*pos*_(*t*), the negative non-coding potential ℰ_*neg*_(*t*), and the discrete translation initiation score ℰ_*tis*_(*t*).

We define a dynamic programming matrix *V*_*t,k*_ storing the maximum accumulated log-probability path up to codon *t* ending in state *k* ∈ {0, 1, 2}. The initialization is set securely to the 5’ region: *V*_0,0_ = 0.0, and *V*_0,1_ = *V*_0,2_ = − ∞. During the single forward pass, the recurrence relations strictly evaluate the biologically permissible state hypotheses:

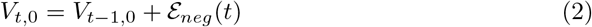

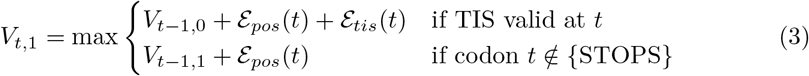

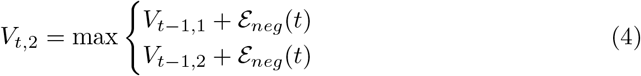

By maintaining an auxiliary pointer matrix to record the optimal incoming state transition, the valid coding structures can be effectively resolved during the traceback phase. In terms of algorithmic complexity, let *N* denote the sequence length in codons and *C* the size of the discrete state space (*C* = 3). While a standard fully-connected Viterbi algorithm operates with a time complexity of *O*(*N* · *C*^2^) per reading frame, the structurally constrained design of NeuroCDS significantly optimizes this computational overhead. Because our formulation enforces a strict, unidirectional biological grammar (5’ UTR →CDS →3’ UTR), the state transition graph is heavily restricted; the algorithm evaluates a maximum of only two permissible incoming transitions per state at any given step. Consequently, the actual time complexity per reading frame collapses to a highly efficient *O*(*N*). Correspondingly, maintaining the dynamic programming and auxiliary traceback matrices necessitates a space complexity of *O*(*N*). This ensures rapid execution suitable for transcriptome-scale data analysis without compromising structural rigor.

### Dynamic Length Normalization and Final Path Selection

A prominent issue in unnormalized Markov models evaluating continuous dense segments is a severe length bias; ultra-long structural ORFs inherently accumulate higher summation scores regardless of average regional confidence. To counter this, NeuroCDS employs a Dynamic Length Normalization mechanism.

Alongside the primary DP matrix, the algorithm dynamically records the accumulated coding sum (𝒮_*CDS*_), the raw number of codons parsed (ℒ_*CDS*_), and the accumulated initiation sum (𝒮_*TIS*_) specific to the trajectory of *S*_1_. Upon completing the forward pass, rather than naively comparing the raw terminal states across different reading frames, NeuroCDS computes a length-balanced final decision score ℱ to select the optimal global transcript structure:

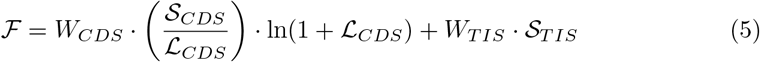

In our implementation, we empirically assign appropriate fusion weights *W*_*CDS*_ and *W*_*TIS*_. This formulation achieves an effective balance: the term (𝒮_*CDS*_*/*ℒ_*CDS*_) guarantees that the region maintains a high average coding density, while the non-linear scaling factor ln(1 + ℒ_*CDS*_) softly rewards structurally longer coding regions without overwhelmingly dominating the TIS evidence (𝒮_*TIS*_). The global reading frame maximizing F is ultimately decoded to yield the robust, final CDS annotation.

## Results

### Performance on Strand-Specific Full-Length Transcripts

The reliable structural annotation of full-length (FL) transcripts provides the fundamental basis for all downstream gene functional analysis and comparative genomics. Evaluating models on intact, strand-specific sequences serves as the primary benchmark to assess the theoretical upper limit of a predictive algorithm. Our comprehensive evaluation across four major eukaryotic clades—Vertebrates, Invertebrates, Plants, and Fungi—reveals a clear performance hierarchy rooted in the underlying algorithmic architectures of the tools (Table 1 and Figure 2A).

**Table 1.**
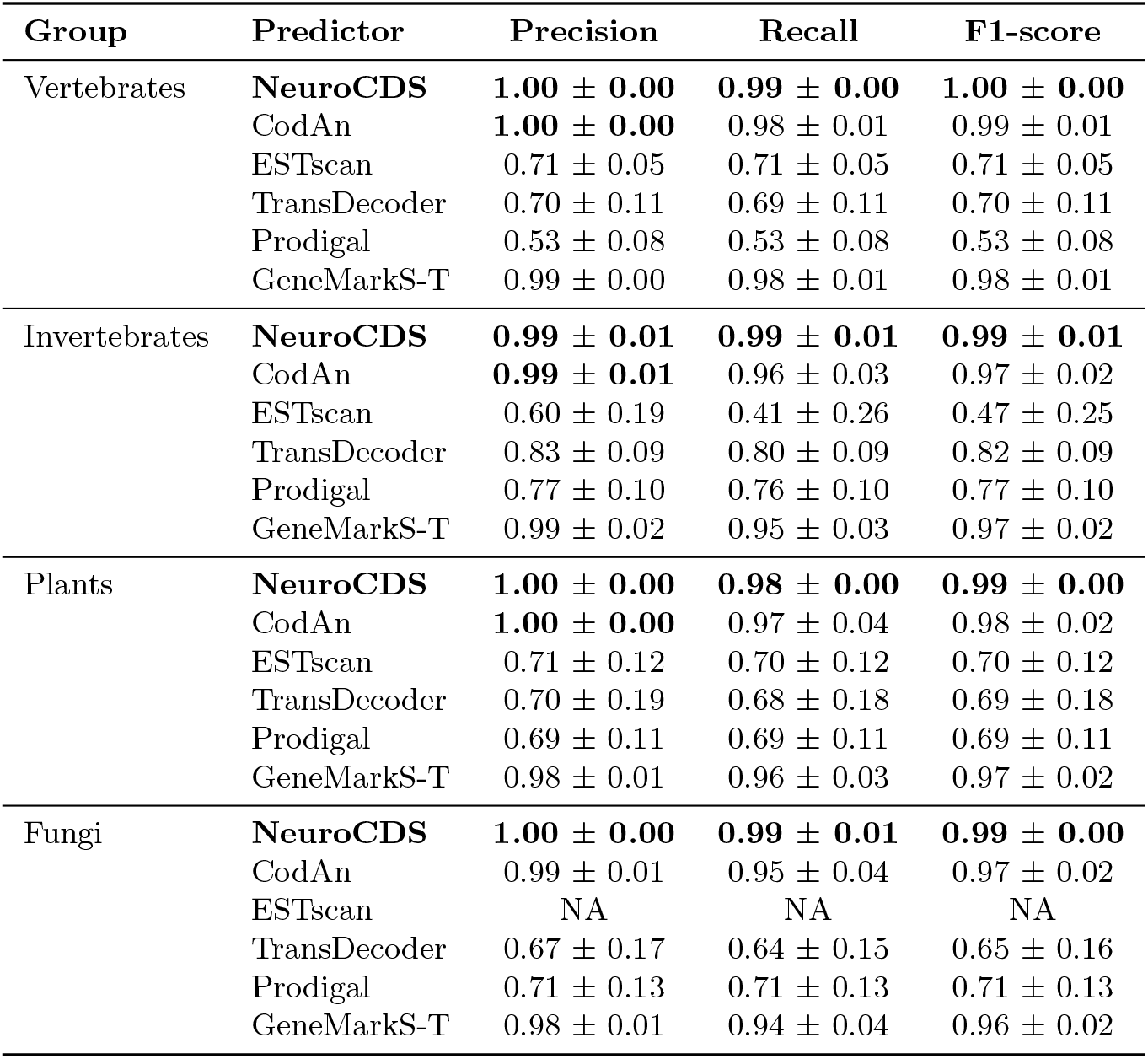
Average and standard deviation of precision, recall, and F1-score obtained by each tool in the strand-specific full transcript sets analyzed. Bold font highlights the highest value for each group. No results are available for ESTscan in fungal transcripts due to the absence of a fungal model.

**Figure 1.**
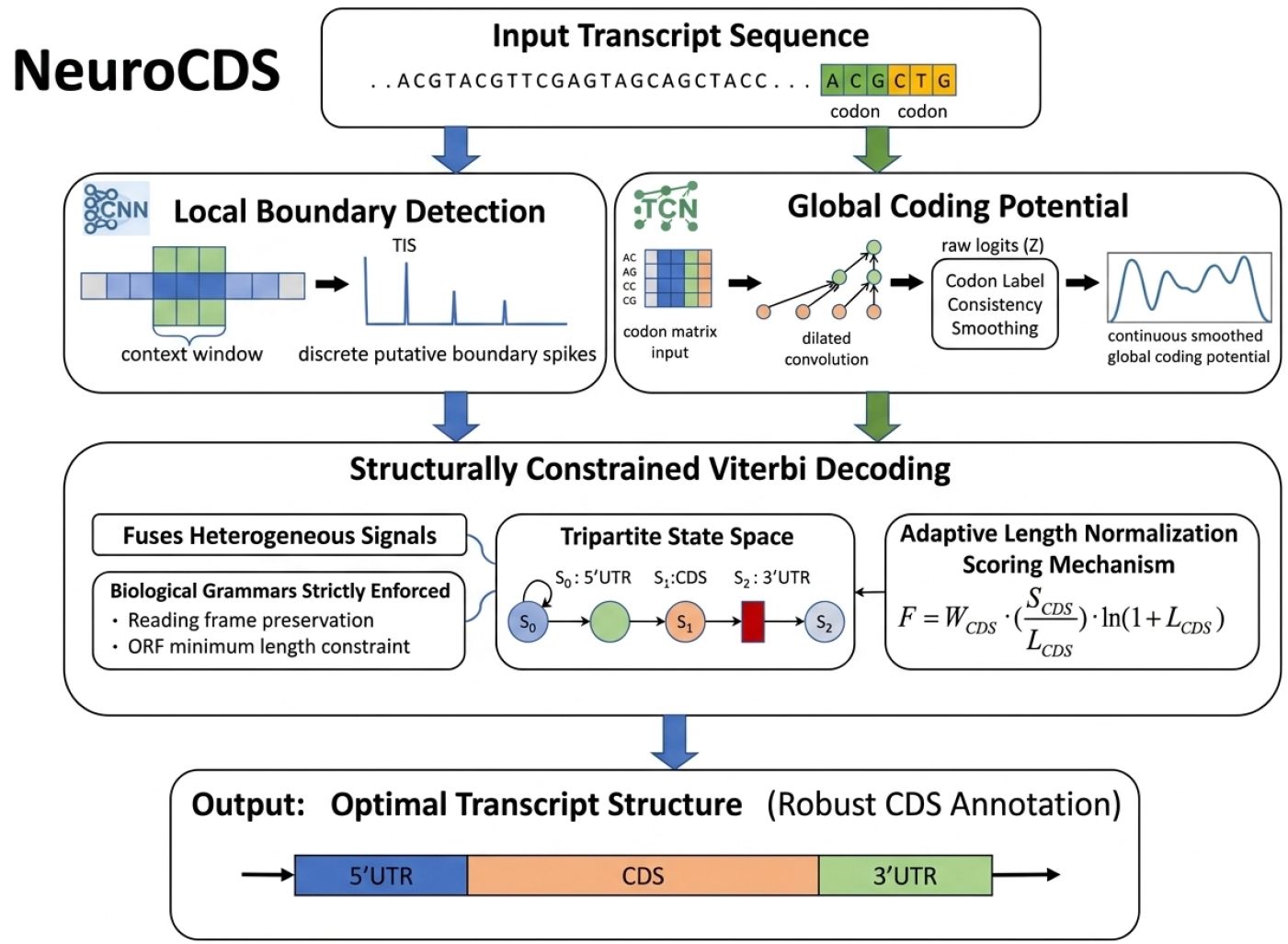
The overall architecture of NeuroCDS. The framework synergistically integrates two neural pathways: a CNN serving as a local sensor to capture discrete structural motifs, and a Temporal Convolutional Network (TCN) acting as a global sensor to evaluate continuous regional coding potential. These heterogeneous representations are subsequently fused via a structurally constrained Viterbi decoding algorithm to reliably delineate the optimal coding sequence.

**Figure 2.**
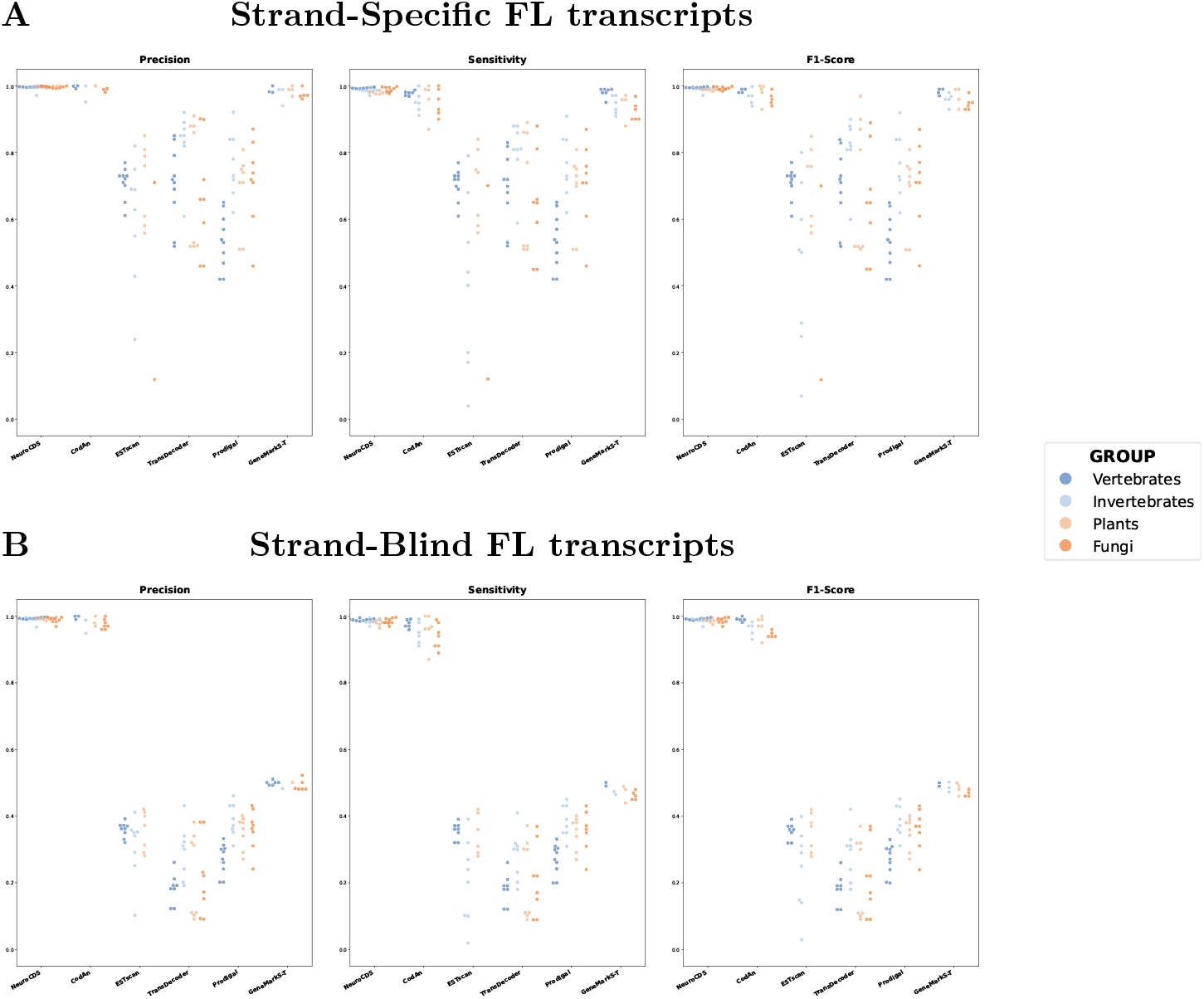
Scatter plot of the full-length (FL) transcript test results. (A) Precision, Sensitivity, and F1-Score obtained by each tool in the strand-specific FL test. (B) Metrics obtained in the strand-blind FL test. Each dot represents a different organism, color-coded by organism group. Results are grouped horizontally by predictor.

While both NeuroCDS and CodAn represent the top tier of well-performing predictors, a detailed analysis of Table 1 reveals a consistent and measurable advantage for NeuroCDS across all evaluated eukaryotic clades. Although CodAn utilizes exquisitely designed Generalized Hidden Markov Models (GHMM) to capture global coding potential, its reliance on fixed statistical transitions inherently caps its sensitivity. In contrast, NeuroCDS consistently outperforms CodAn, particularly in Recall (Sensitivity) and the comprehensive F1-score. As demonstrated in Table 1, NeuroCDS achieves a high Recall of 0.98 to 0.99 across all clades, whereas CodAn’s Recall notably drops to 0.95 in Fungi and 0.96 in Invertebrates. Beyond absolute metric values, the standard deviations highlight the exceptional stability of NeuroCDS. While CodAn and GeneMarkS-T exhibit noticeable performance fluctuations (with standard deviations up to ± 0.04) when encountering diverse codon usage biases across different kingdoms, NeuroCDS uniformly maintains near-zero standard deviations (± 0.00 to ± 0.01) across Precision, Recall, and F1-score. This minimal dispersion mathematically proves the superiority of our algorithmic design. By replacing static Markov transition tables with high-dimensional neural representations, NeuroCDS not only preserves the structural rigor of GHMMs but also excels at extracting robust, non-linear sequence features that remain highly stable regardless of organism-specific sequence complexity. The synergistic integration of multi-scale features—a CNN for Translation Initiation Site (TIS) localization and a TCN for global regional coding evaluation—enables NeuroCDS to significantly reduce false negatives while universally maintaining a robust Precision of 0.99 to 1.00. GeneMarkS-T follows closely behind this leading group, but as previously noted, its traditional hexamer-based statistical models are less expressive than deep learning architectures, resulting in slightly higher performance variance.

In contrast, tools relying on heuristics or non-specialized models constitute the other baseline tools with substantial performance fluctuations across species. TransDecoder, which relies on identifying the longest Open Reading Frame (ORF) and scoring via likelihood ratio tests, shows F1-scores primarily clustered between 0.60 and 0.85. This limitation arises because the longest ORF does not always coincide with the functional coding region, and the lack of a joint structural inference mechanism leads to the misidentification of UTR segments. Prodigal [18] suffers from an evident domain mismatch; originally optimized for prokaryotic gene finding, it lacks the sensitivity to distinguish eukaryotic-specific signals like the Kozak consensus, resulting in F1-scores as low as 0.53 in vertebrate datasets. Similarly, ESTscan exhibits the most significant dispersion, with performance exhibiting significant fluctuations in several organisms. This reflects the constrained adaptability of its older scoring matrices and traditional Markov models which are less suited for the complex, non-linear sequence patterns found across modern eukaryotic transcriptomes.

Overall, the high metric accuracy combined with the unmatched statistical consistency (near-zero standard deviations) of NeuroCDS across all clades underscores the advantage of coupling deep learning representations with a mathematically rigorous Segmental Viterbi decoder. This hybrid approach ensures that NeuroCDS provides a more robust and broadly adaptable solution for full-length transcript annotation than both traditional heuristic tools and purely statistical methods like GHMMs.

### Robust Identification in Strand-Blind Scenarios

In transcriptomics, *de novo* assembly frequently generates contigs without strand orientation, particularly when using non-directional RNA-Seq libraries. A robust prediction model must reliably parse coding regions across all six possible reading frames (three per strand) to prevent massive data loss in unoriented assemblies. Our evaluation using strand-blind full-length datasets (Table 2 and Figure 2B) demonstrates a significant divergence in performance, highlighting the necessity of global structural modeling in mitigating orientation ambiguity.

**Table 2.** Average and standard deviation of precision, recall, and F1-score obtained by each tool in the strand-blind full transcript sets analyzed. Bold font highlights the highest value for each group. No results are available for ESTscan in fungal transcripts due to the absence of a fungal model.

| Group | Predictor | Precision | Recall | F1-score |
| --- | --- | --- | --- | --- |
| Vertebrates | **NeuroCDS** | **0.99 ± 0.00** | **0.99 ± 0.00** | **0.99 ± 0.00** |
|  | CodAn | **0.99 ± 0.00** | 0.98 ± 0.01 | **0.99 ± 0.01** |
|  | ESTscan | 0.36 ± 0.02 | 0.36 ± 0.02 | 0.88 ± 0.13 |
|  | TransDecoder | 0.18 ± 0.04 | 0.18 ± 0.04 | 0.93 ± 0.05 |
|  | Prodigal | 0.27 ± 0.05 | 0.27 ± 0.05 | 0.93 ± 0.02 |
|  | GeneMarkS-T | 0.50 ± 0.00 | 0.49 ± 0.00 | 0.49 ± 0.00 |
| Invertebrates | **NeuroCDS** | **0.99 ± 0.01** | **0.98 ± 0.01** | **0.99 ± 0.01** |
|  | CodAn | **0.99 ± 0.02** | 0.95 ± 0.03 | 0.97 ± 0.02 |
|  | ESTscan | 0.31 ± 0.10 | 0.21 ± 0.13 | 0.24 ± 0.12 |
|  | TransDecoder | 0.29 ± 0.08 | 0.28 ± 0.07 | 0.29 ± 0.08 |
|  | Prodigal | 0.39 ± 0.05 | 0.39 ± 0.05 | 0.39 ± 0.05 |
|  | GeneMarkS-T | 0.49 ± 0.01 | 0.48 ± 0.01 | 0.48 ± 0.01 |
| Plants | **NeuroCDS** | **0.99 ± 0.00** | **0.98 ± 0.01** | **0.99 ± 0.01** |
|  | CodAn | **0.99 ± 0.01** | 0.96 ± 0.04 | 0.98 ± 0.02 |
|  | ESTscan | 0.35 ± 0.06 | 0.35 ± 0.06 | 0.35 ± 0.06 |
|  | TransDecoder | 0.22 ± 0.13 | 0.21 ± 0.12 | 0.21 ± 0.12 |
|  | Prodigal | 0.35 ± 0.05 | 0.35 ± 0.05 | 0.35 ± 0.05 |
|  | GeneMarkS-T | 0.49 ± 0.01 | 0.48 ± 0.02 | 0.49 ± 0.01 |
| Fungi | **NeuroCDS** | **0.99 ± 0.01** | **0.99 ± 0.01** | **0.99 ± 0.01** |
|  | CodAn | 0.98 ± 0.01 | 0.94 ± 0.04 | 0.96 ± 0.02 |
|  | ESTscan | NA | NA | NA |
|  | TransDecoder | 0.21 ± 0.11 | 0.20 ± 0.10 | 0.21 ± 0.11 |
|  | Prodigal | 0.36 ± 0.06 | 0.35 ± 0.06 | 0.36 ± 0.06 |
|  | GeneMarkS-T | 0.49 ± 0.01 | 0.47 ± 0.02 | 0.48 ± 0.01 |

While both NeuroCDS and CodAn demonstrate resilience in this challenging scenario, a close examination reveals that NeuroCDS maintains a distinct superiority in both metric magnitude and statistical stability. Although CodAn utilizes its GHMM architecture to filter non-functional strands, its sensitivity experiences a noticeable decline (e.g., Recall dropping to 0.94 in Fungi) accompanied by increased variance (standard deviations up to ± 0.04). In stark contrast, NeuroCDS consistently dominates with Recall scores of 0.98 to 0.99 and near-zero standard deviations (± 0.00 to ± 0.01) across all taxonomic groups. This superior stability stems from our innovative fusion strategy. The Segmental Viterbi algorithm in NeuroCDS independently evaluates the joint probability of continuous coding signals (via TCN) and local sequence motifs (via CNN) for each of the six reading frames. By maximizing the global path probability across this high-dimensional neural representation space, the framework organically and decisively suppresses the deceptive, fragmented ORF-like shadows typically found on the reverse-complement strand—a task where traditional Markovian transition probabilities show vulnerability.

The advantage of our approach becomes even more pronounced when compared to baseline tools—including ESTscan, TransDecoder, and Prodigal—which experienced catastrophic performance degradation. As illustrated in Figure 2B, their Precision and Recall metrics collapsed to the 0.20–0.40 range for most clades. This severe failure can be attributed to their lack of robust strand-invariant modeling. TransDecoder and Prodigal rely heavily on directional heuristics, such as prioritizing the single longest ORF. When confronted with both strands simultaneously, these tools completely fail to distinguish the true CDS from pseudo-signals on the reverse strand, frequently outputting multiple overlapping or entirely false predictions. GeneMarkS-T also exhibited significant degradation, with F1-scores dropping to approximately 0.48–0.49. While it attempts to model both strands, its traditional statistical scoring lacks the necessary high-dimensional discriminative power to reliably separate the true signal from complex reverse-complement noise.

These results emphatically prove that local motif detection or simple ORF-length heuristics are grossly insufficient for unoriented transcriptome annotation. The ability of NeuroCDS to maintain uncompromised accuracy and near-zero variance in strand-blind scenarios confirms that integrating synergistic deep learning representations with a rigorous Viterbi decoder provides a definitive advantage. By treating the entire six-frame landscape as a holistic sequence parsing problem driven by deep features, NeuroCDS ensures that the final annotation remains remarkably robust even when experimental orientation data is entirely missing.

### Evaluation on Complex Ribo-seq Validated Datasets

While computational benchmarks against reference genomes verify a model’s theoretical accuracy, real-world biological sequences exhibit greater complexity, including alternative transcript isoforms, non-coding noise, and dataset-specific sequencing biases. Ribosome profiling (Ribo-seq) provides direct experimental evidence of active translation, making it a valuable dataset for assessing a methodology’s capability to identify functional coding regions *in vivo*. Our evaluation across multiple species and experimental sources (Table 3 and Figure 3) compares the performance of NeuroCDS with existing annotation models.

**Table 3.** Precision, recall, and F1-score for the prediction in datasets validated by ribo-seq experiments. Bold font highlights the highest value for each dataset. The ‘Size’ column refers to the number of transcripts in the dataset.

| Dataset | Size | Predictor | Precision | Recall | F1-score |
| --- | --- | --- | --- | --- | --- |
| *H. sapiens* (Lim et al., 2018) | 14193 | **NeuroCDS** | **0.96** | **0.98** | **0.97** |
|  |  | CodAn | 0.81 | 0.75 | 0.78 |
|  |  | ESTScan | 0.23 | 0.22 | 0.23 |
|  |  | TransDecoder | 0.37 | 0.35 | 0.36 |
|  |  | Prodigal | 0.27 | 0.27 | 0.27 |
|  |  | GeneMarkS-T | 0.67 | 0.63 | 0.65 |
| *H. sapiens* (Lee et al., 2012) | 5727 | **NeuroCDS** | **1.00** | **0.99** | **1.00** |
|  |  | CodAn | 0.99 | 0.97 | 0.98 |
|  |  | ESTScan | 0.68 | 0.68 | 0.68 |
|  |  | TransDecoder | 0.60 | 0.59 | 0.60 |
|  |  | Prodigal | 0.49 | 0.49 | 0.49 |
|  |  | GeneMarkS-T | 0.98 | 0.96 | 0.97 |
| *M. musculus* (Lim et al., 2018) | 20326 | **NeuroCDS** | **0.93** | **0.97** | **0.95** |
|  |  | CodAn | 0.85 | 0.83 | 0.84 |
|  |  | ESTScan | 0.30 | 0.30 | 0.30 |
|  |  | TransDecoder | 0.39 | 0.38 | 0.39 |
|  |  | Prodigal | 0.29 | 0.29 | 0.29 |
|  |  | GeneMarkS-T | 0.70 | 0.68 | 0.69 |
| *M. musculus* (Lee et al., 2012) | 2701 | **NeuroCDS** | **1.00** | **1.00** | **1.00** |
|  |  | CodAn | **1.00** | 0.98 | 0.99 |
|  |  | ESTScan | 0.74 | 0.74 | 0.74 |
|  |  | TransDecoder | 0.67 | 0.65 | 0.66 |
|  |  | Prodigal | 0.52 | 0.52 | 0.52 |
|  |  | GeneMarkS-T | 0.97 | 0.95 | 0.97 |
| *D. rerio* (Lim et al., 2018) | 13954 | **NeuroCDS** | **1.00** | **0.99** | **1.00** |
|  |  | CodAn | 0.86 | 0.85 | 0.86 |
|  |  | ESTScan | 0.44 | 0.44 | 0.44 |
|  |  | TransDecoder | 0.67 | 0.65 | 0.66 |
|  |  | Prodigal | 0.47 | 0.47 | 0.47 |
|  |  | GeneMarkS-T | 0.80 | 0.78 | 0.79 |
| *D. melanogaster* (Lim et al., 2018) | 13653 | **NeuroCDS** | **0.99** | **0.98** | **0.99** |
|  |  | CodAn | 0.90 | 0.90 | 0.90 |
|  |  | ESTScan | 0.68 | 0.66 | 0.67 |
|  |  | TransDecoder | 0.57 | 0.56 | 0.56 |
|  |  | Prodigal | 0.56 | 0.56 | 0.56 |
|  |  | GeneMarkS-T | 0.86 | 0.83 | 0.85 |
| *A. thaliana* (Lim et al., 2018) | 6947 | **NeuroCDS** | **1.00** | **0.98** | **0.99** |
|  |  | CodAn | 0.87 | 0.85 | 0.86 |
|  |  | ESTScan | 0.59 | 0.58 | 0.58 |
|  |  | TransDecoder | 0.67 | 0.65 | 0.66 |
|  |  | Prodigal | 0.56 | 0.56 | 0.56 |
|  |  | GeneMarkS-T | 0.82 | 0.79 | 0.80 |

**Figure 3.**
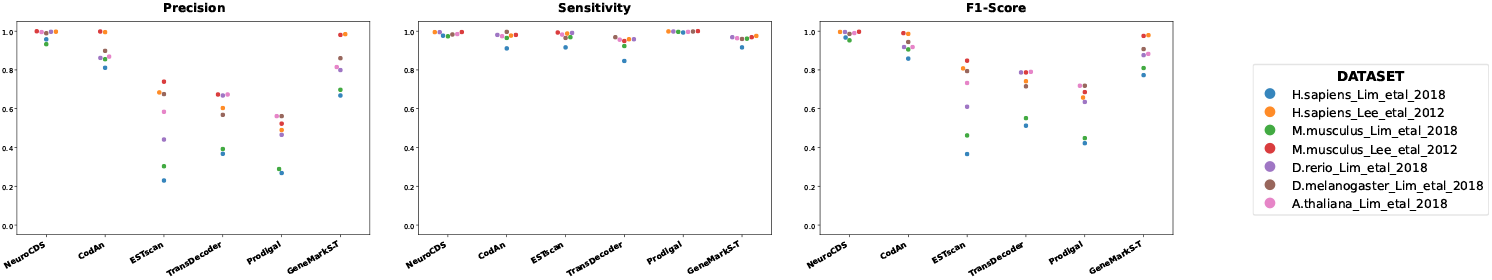
Scatter plot of the precision, sensitivity, and F1-score obtained by each tool in the Ribo-seq experimentally validated datasets. Each dot represents a different dataset, color-coded by dataset name.

As shown in Table 3, NeuroCDS consistently maintained F1-scores between 0.95 and 1.00 across all seven Ribo-seq datasets. Performance variations among the evaluated tools were particularly noticeable between the different dataset sources. On the Lee et al. (2012) datasets, multiple tools, including NeuroCDS, CodAn, and GeneMarkS-T, performed competitively, all achieving F1-scores of 0.97 or higher for both *H. sapiens* and *M. musculus*.

However, on the Lim et al. (2018) datasets, which present a more challenging sequence environment, traditional statistical models exhibited decreased sensitivity. For instance, CodAn’s F1-score was 0.78 for *H. sapiens* and 0.84 for *M. musculus*, while GeneMarkS-T achieved 0.65 and 0.69, respectively. In contrast, NeuroCDS sustained F1-scores of 0.97 and 0.95 on these same datasets. This divergence highlights a limitation of static probabilistic models: predefined Markov transition probabilities can be sensitive to variations in experimental noise or deviations from standard emission profiles. The stability of NeuroCDS across different experimental conditions can be attributed to its dual-branch neural architecture. By requiring joint validation from both localized sequence motifs (via the CNN) and continuous regional signals (via the TCN), the model effectively mitigates the impact of transient sequencing anomalies.

Other baseline predictors, including TransDecoder, Prodigal, and ESTscan, yielded substantially lower F1-scores (frequently below 0.40) across the analyzed Ribo-seq datasets. This performance gap indicates that naive longest-ORF heuristics or simpler sequence scanning tools are generally insufficient for distinguishing genuine *in vivo* translation sites from non-functional, ORF-like segments in complex transcriptomic data.

### Robustness on Fragmented Assemblies

A ubiquitous limitation of *de novo* transcriptome assembly algorithms is the frequent generation of incomplete contigs due to low sequencing depth or complex repetitive regions. These partial transcripts may lack the 5’ end, the 3’ end, or both. Evaluating predictive models on these fragmented assemblies is crucial for assessing their practical utility in real-world bioinformatics pipelines. Our analysis of three distinct fragmentation scenarios (Figure 4) reveals how different algorithmic strategies cope with the loss of key sequence signatures.

**Figure 4.**
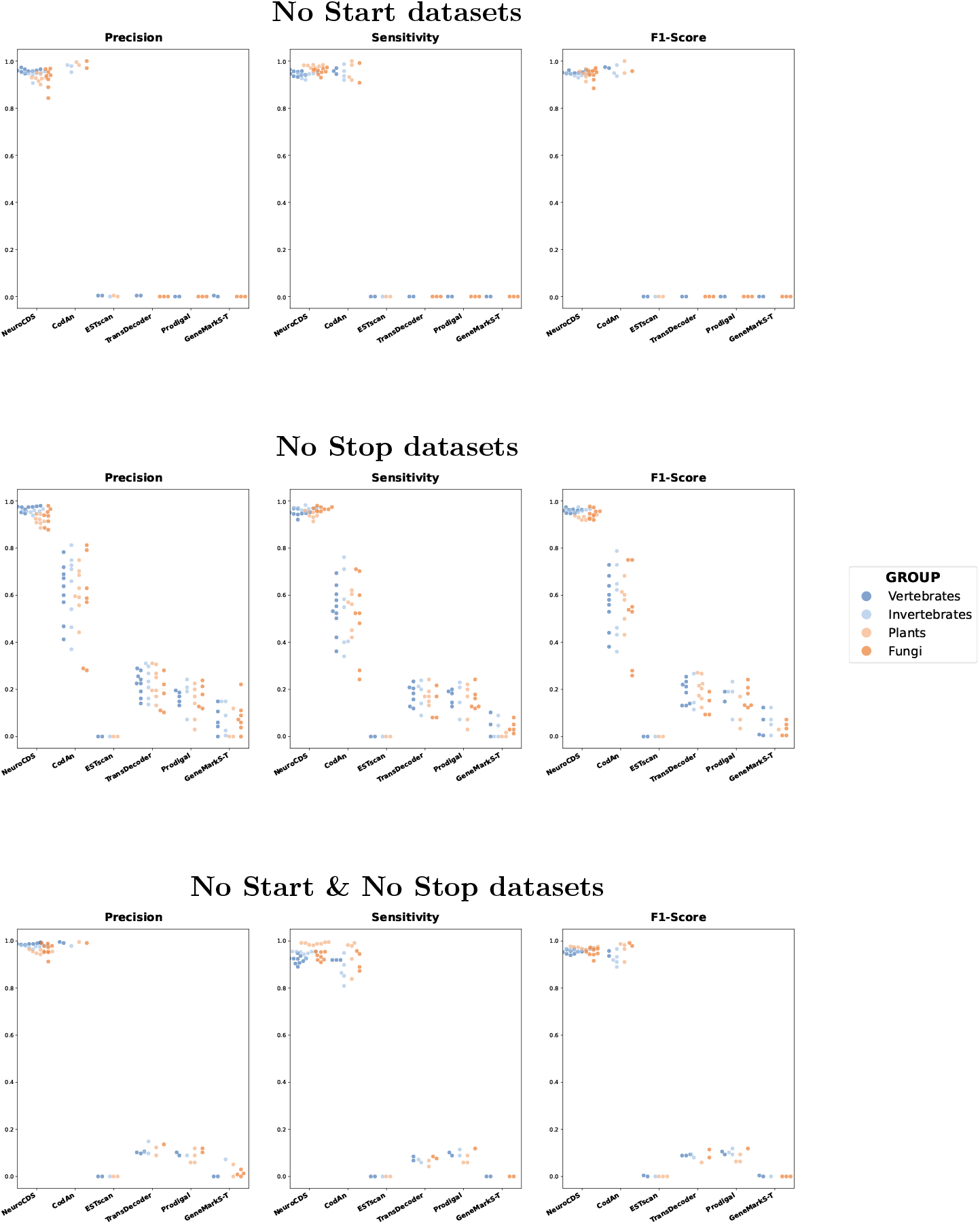
Scatter plot of the partial transcript test results. The plots show the Precision, Sensitivity, and F1-Score obtained by each tool on ‘No Start’, ‘No Stop’, and ‘No Start & No Stop’ tests performed on all analyzed species. Each dot represents a different organism, color-coded by organism group.

#### 5’-Truncated Fragments (NoStart)

For contigs missing upstream sequences, both CodAn and NeuroCDS demonstrated competitive performance, albeit with distinct metric profiles. CodAn achieved high Precision (approaching 1.0), likely due to its GHMM architecture dynamically adjusting initial state probabilities to accommodate missing initiation contexts. Conversely, NeuroCDS exhibited higher Recall (predominantly between 0.90 and 0.98), though with slightly increased variance in Precision compared to full-length evaluations. From an algorithmic perspective, the absence of the discrete TIS signal introduces ambiguity in identifying the exact start site; however, the continuous regional representations generated by the TCN branch compensate by anchoring the prediction on the sustained coding signature, thereby minimizing false negatives in internal coding regions.

#### 3’-Truncated Fragments (NoStop)

The robustness of NeuroCDS is most prominent in 3’-truncation scenarios, where downstream termination signals are absent. As illustrated in Figure 4, NeuroCDS maintained high F1-scores, with Precision and Recall universally exceeding 0.90 across diverse clades. This resilience is a direct result of the synergistic architecture of the framework. Notably, the maintained high Precision is likely attributable to the TIS CNN module, which acts as a reliable 5’ structural anchor. By accurately capturing the translation initiation context, the CNN provides a high-confidence starting state that restricts the Viterbi decoder from falsely initiating on spurious ORFs. Furthermore, even without a stop codon signal, the decoder can stably evaluate the length of the coding region based on the sustained regional coding potential from the TCN and the dynamic length normalization mechanism. In contrast, traditional statistical models like CodAn exhibited broader performance fluctuations in this category, with F1-scores for many species falling into the 0.20–0.80 range. This suggests that the absence of a termination state transition can occasionally derail Markov-based paths that rigidly expect a structured exit from the coding state.

#### Internal Fragments (NoStartNoStop)

Sequences lacking both canonical boundaries represent the most information-poor scenario, heavily penalizing methods reliant on exact motif matching. In this category, NeuroCDS maintained a stable performance profile, with F1-scores consistently exceeding 0.95 across most evaluated clades. This stability highlights the advantage of framing CDS annotation as a continuous structural parsing problem: the Segmental Viterbi algorithm can still resolve the optimal reading frame based solely on the TCN’s codon usage evaluations. While CodAn retained high Precision, its Recall exhibited noticeably higher variance than NeuroCDS. Other baseline tools (TransDecoder, Prodigal, and ESTscan) demonstrated severe performance degradation, primarily because their heuristic scoring and static models depend heavily on canonical start/stop codons or intact sequence lengths.

Overall, these evaluations confirm that the synergistic integration of discrete local motifs and continuous global representations provides substantial robustness against assembly artifacts. By leveraging regional sequence context to compensate for missing boundary signals, NeuroCDS facilitates the reliable recovery of functional coding sequences from the heavily fragmented transcriptomes typical of non-model organisms.

### Impact of Sequence Length Degradation

The physical length of an assembled contig intrinsically dictates the volume of biological context available for predictive algorithms. Extremely short sequences, or “micro-fragments,” pose a dual challenge: they often lack complete canonical motifs for localized sensors and fail to provide sufficient length for stable long-range statistical patterns. To systematically quantify the limits of NeuroCDS, we subjected human and mouse datasets to progressive stochastic truncation, generating three distinct length bins: 500–1000 nt, 200–500 nt, and 50–200 nt.

Crucially, to ensure a rigorous evaluation of these truncated segments, the ground-truth reading frame was not naively inherited from the parental full-length transcript. Instead, we implemented a precise coordinate-mapping protocol: the true reading frame of each fragment was determined by calculating the base-0 positional offset relative to the original sequence, with the frame assigned via a modulo 3 synchronization logic. This meticulous approach established a rigorous and biologically consistent baseline for assessing fragmented assemblies.

The experimental results (Table 4) demonstrate that NeuroCDS maintains good robustness on segments exceeding 200 nt, with all metrics remaining within a high-utility range. Predictably, performance exhibited a moderate decline on the extreme 50–200 nt micro-fragments, with Specificity in humans adjusting to 90.26% and Sensitivity in mouse to 89.45%. This trend provides transparent mechanistic insights into the framework’s behavior: extreme truncation limits the spatial context required for the CNN branch to identify TIS signals, while simultaneously restricting the TCN’s ability to accumulate sufficient cross-codon dependencies for global regional evaluation.

**Table 4.** Impact of sequence length degradation on prediction performance using human and mouse datasets. Metrics evaluated include Sensitivity (SN), Specificity (SP), and Area Under the ROC Curve (auROC).

| Length | human |  |  | mouse |  |  |
| --- | --- | --- | --- | --- | --- | --- |
|  | SN(%) | SP(%) | auROC | SN(%) | SP(%) | auROC |
| 500–1000 nt | 98.57 | 99.18 | 0.9988 | 98.79 | 99.12 | 0.9988 |
| 200–500 nt | 96.45 | 98.74 | 0.9968 | 98.07 | 97.81 | 0.9973 |
| 50–200 nt | 95.86 | 90.26 | 0.9821 | 89.45 | 96.13 | 0.9828 |

However, despite this contextual scarcity, the model’s auROC remained relatively high even in the shortest bin. This indicates that while stable identification becomes more difficult as sequences approach the limits of the neural receptive fields, the core architecture of NeuroCDS retains a fundamental capacity to distinguish functional coding signatures from non-coding noise. This graceful degradation confirms the framework’s resilience in the face of the heavy fragmentation characteristic of real-world *de novo* transcriptomes.

### Specificity and False-Positive Suppression

A critical requirement for transcriptome annotation is the ability to reject non-coding artifacts. Raw assemblies frequently contain abundant non-coding elements, including 3’ UTR fragments and long non-coding RNAs (ncRNAs). Falsely annotating these sequences as valid coding regions severely confounds downstream proteomic databases and functional genomic studies. We evaluated the models on dedicated 3’UTR and ncRNA datasets to assess their Specificity and false-discovery resilience (Figure 5).

**Figure 5.**
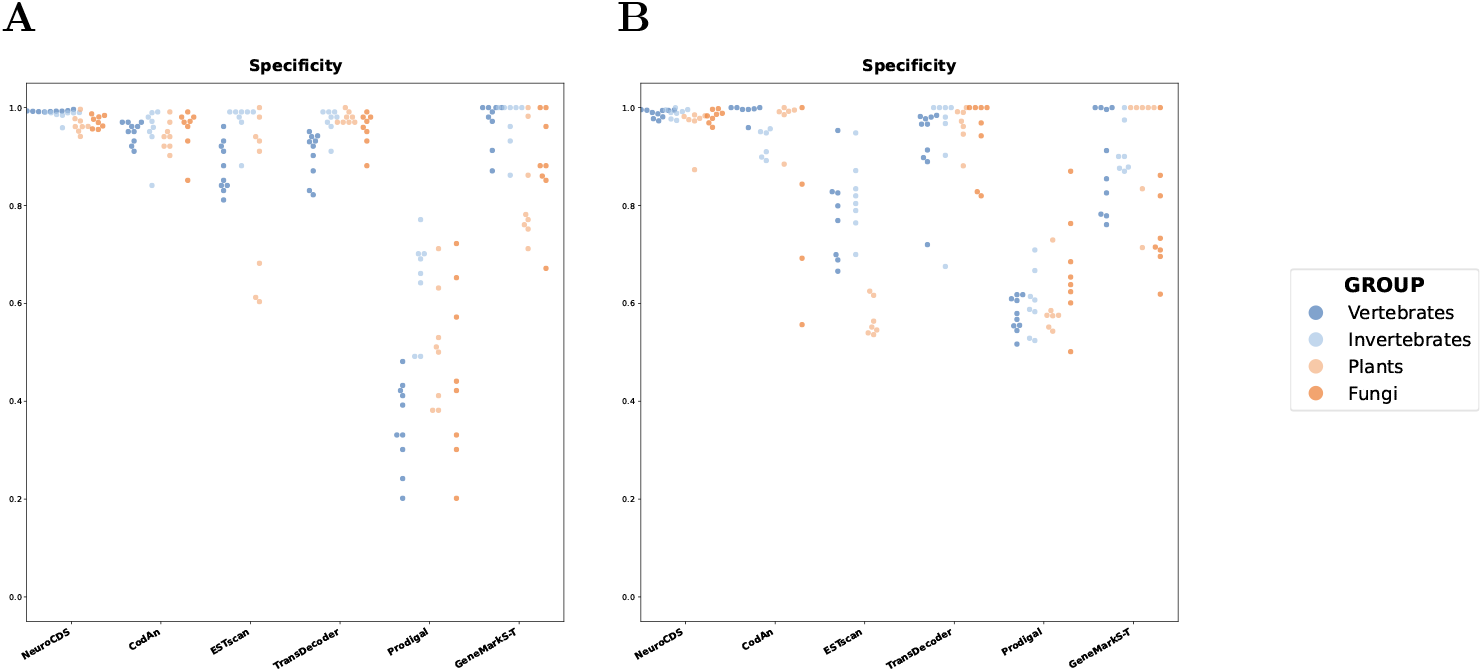
Scatter plot of Specificity obtained by each tool in the prediction. We used two negative datasets: (A) 3’UTR region datasets and (B) ncRNA datasets. Each dot represents a different organism, color-coded by organism group.

As illustrated in Figure 5, NeuroCDS consistently maintained Specificity scores approaching 1.0 across both datasets, exhibiting minimal variance across different taxonomic groups. This stringent false-positive suppression is attributable to the joint inference requirement of its dual-branch architecture. Because the Viterbi decoder strictly integrates both the discrete spatial motif recognition from the CNN and the continuous codon usage evaluation from the TCN, a non-coding artifact must simultaneously present canonical translation initiation signals and sustained triplet-based coding potential to be falsely classified. This structural constraint effectively filters out deceptive pseudo-ORFs that may possess localized similarities to true genes but lack global compositional consistency.

In contrast, traditional probabilistic models exhibited distinct vulnerabilities, particularly when transitioning from 3’ UTRs to more structurally complex ncRNAs. While CodAn maintained relatively stable Specificity on the 3’ UTR dataset (Figure 5A), its performance on the ncRNA dataset (Figure 5B) revealed a broader dispersion, with scores for several organisms experiencing noticeable declines. This pattern highlights a known limitation of static Markov-based models: complex ncRNAs frequently harbor pseudo-ORFs that inadvertently mirror the hexamer frequencies or compositional emission profiles of genuine genes, causing rigid probabilistic transition matrices to misclassify them. GeneMarkS-T exhibited a similar susceptibility to ncRNA artifacts, showing wide inter-species variance.

The heuristic and non-specialized baseline tools—ESTscan, TransDecoder, and Prodigal—demonstrated substantial Specificity degradation and high cross-species variance, with numerous data points scattering well below the 0.60 threshold.TransDecoder’s reliance on a longest-ORF heuristic makes it particularly prone to misidentifying coincidental, long open reading frames within unconstrained non-coding regions as functional CDS. Similarly, Prodigal’s primary optimization for prokaryotic gene structures renders it poorly equipped to differentiate eukaryotic non-coding architectures from true transcripts. Overall, the data indicates that integrating multi-scale neural feature extraction with structural Viterbi decoding provides a measurable advantage in minimizing false discoveries within the noise-rich environments characteristic of *de novo* transcriptomics.

### Overall Evaluation on Combined Datasets

To systematically evaluate model robustness across highly heterogeneous sequencing scenarios, we aggregated all previously defined sequence types—including full-length transcripts, diverse partial fragments, and non-coding artifacts—into a combined evaluation pool. This mixed dataset simulates the unpredictable and complex landscape of raw *de novo* transcriptome assemblies. The results (Figure 6) highlight distinct behavioral divergences among the evaluated annotation methodologies.

**Figure 6.**
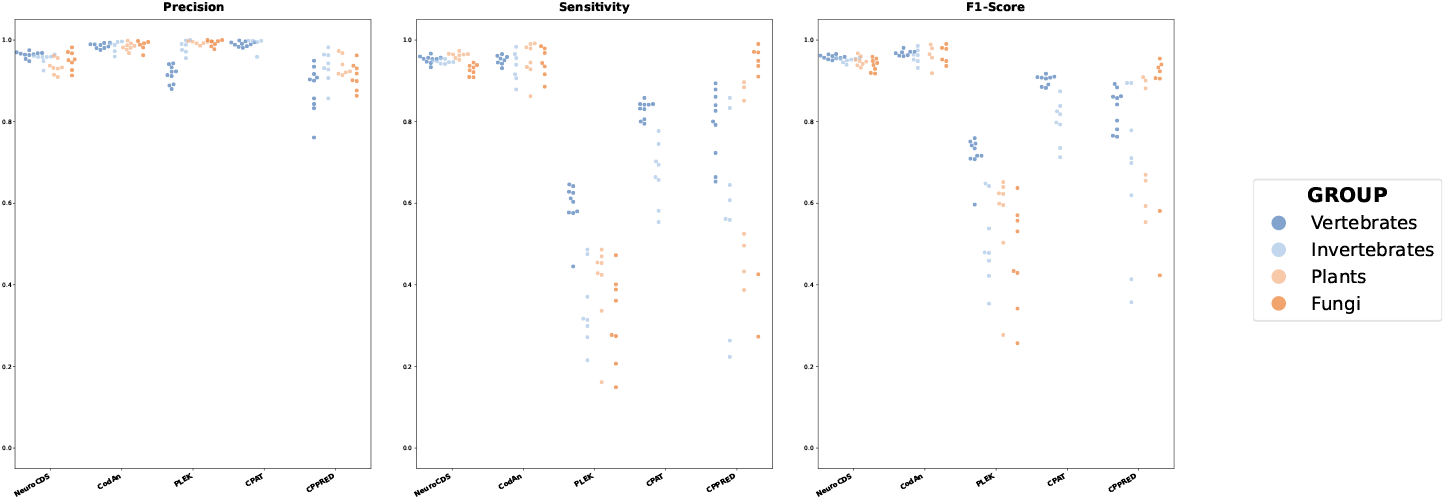
Scatter plot for Precision, Sensitivity, and F1-score obtained by evaluating all designed datasets combined. Each dot represents a different organism, color-coded by organism group.

As depicted in the F1-score distributions, NeuroCDS and CodAn establish the top-performing tier. NeuroCDS exhibits the highest clustering density at the upper limit of the F1-score range, indicating highly consistent structural parsing regardless of species or sequence integrity. In contrast, alignment-free or heuristic-based baseline tools, such as PLEK [19], CPAT [9], and CPPRED [20], exhibit severe performance dispersion. Their F1-scores scatter widely between 0.20 and 0.90, demonstrating that algorithms lacking robust structural grammar constraints are highly vulnerable to sequence fragmentation and non-coding noise.

Deconstructing the F1-scores into Precision and Sensitivity reveals the underlying algorithmic trade-offs. CodAn demonstrates exceptionally high and tightly clustered Precision, consistently remaining above the 0.95 threshold. This metric profile reflects the inherent stringency of its GHMM architecture: pre-calculated, static state-transition penalties aggressively filter out low-confidence or heavily truncated segments, effectively minimizing false-positive assignments. CPAT similarly exhibits clustered high Precision, but the accompanying Sensitivity plot reveals that this comes at the cost of catastrophic false-negative rates on fragmented sequences.

Conversely, NeuroCDS demonstrates a clear algorithmic prioritization of Sensitivity. While CodAn’s strict Markov paths lead to increased false negatives on atypical sequences (visible as a broader downward spread in its Sensitivity plot), NeuroCDS maintains tightly clustered Sensitivity scores approaching 1.0. However, this robust Recall is accompanied by slightly more distributed Precision scores (predominantly within the 0.90–0.98 range). This dynamic illustrates the intrinsic mechanism of the neural framework: the TCN’s powerful capacity to recognize continuous regional coding potentials allows it to reliably salvage fragmented genes, but in ultra-high-noise environments, this highly sensitive joint probability space may occasionally admit highly gene-like pseudo-ORFs.

Ultimately, the superior stability and tight clustering of NeuroCDS in the comprehensive F1-score evaluation confirm that its synergistic neural integration strategy yields a highly balanced and reliable annotation. By effectively mitigating the extreme sensitivity drops seen in static models when confronted with structural degradation, NeuroCDS provides a highly practical solution for the full spectrum of transcriptome annotation challenges.

### Cross-Species Generalization Potential

Eukaryotic organisms exhibit distinct genomic characteristics, including varying GC contents and divergent codon usage biases. However, the fundamental biological logic of amino acid coding and translation initiation remains largely conserved across clades. To evaluate whether the neural representations learned by NeuroCDS capture these generalized evolutionary principles rather than overfitting to species-specific patterns, we assessed its cross-taxonomic generalization potential by applying the vertebrate-trained model directly to the distantly related plant dataset, *Arabidopsis thaliana* (Table 5).

**Table 5.** Cross-species generalization performance. The table presents the Precision, Recall, F1-score, and Specificity obtained when evaluating the *A. thaliana* (plant) dataset using the pre-trained vertebrate model.

| Sequence Type | Precision | Recall | F1-score |
| --- | --- | --- | --- |
| Strand-specific full | 0.99 | 0.98 | 0.99 |
| NoStart | 0.94 | 0.90 | 0.92 |
| NoStop | 0.92 | 0.89 | 0.90 |
| NoStartNoStop | 0.98 | 0.82 | 0.89 |
| Sequence Type | Specificity |  |  |
| 3'UTR | 0.98 |  |  |
| ncRNA | 0.97 |  |  |

On intact, strand-specific full-length transcripts, the heterologous model achieved an F1-score of 0.99 (Precision: 0.99, Recall: 0.98), indicating that the core translational features extracted by the dual-branch network are highly transferable. When evaluating fragmented assemblies, the model demonstrated varying degrees of performance adjustment. For 5’-truncated (NoStart) and 3’-truncated (NoStop) fragments, the F1-scores were 0.92 and 0.90, respectively. In the most structurally deficient scenario (NoStartNoStop), Precision remained high at 0.98, while Recall adjusted to 0.82. This reduction in Recall highlights the increased difficulty of stably delineating coding regions when discrete spatial motifs are absent and cross-species codon usage deviations accumulate across the continuous TCN evaluation window. Despite this, the vertebrate model successfully maintained stringent false-positive suppression on plant non-coding datasets, recording Specificities of 0.98 on 3’ UTRs and 0.97 on ncRNAs.

These empirical results suggest that the integration of multi-scale neural features with structurally constrained Viterbi decoding leverages conserved biological grammar rather than rigid sequence homology. From a practical bioinformatics perspective, this generalization capacity enables the deployment of pre-trained NeuroCDS models as a viable zero-shot diagnostic tool for annotating *de novo* transcriptomes in non-model organisms or orphan species lacking sufficient high-quality data for species-specific training.

### Computational Efficiency

In the practical deployment of bioinformatics software, computational efficiency is a critical metric alongside predictive accuracy. To evaluate the throughput of NeuroCDS, we recorded its execution time on a comprehensive benchmark pool comprising 82,501 complex transcripts derived from the seven Ribo-seq validated datasets. All benchmarks were conducted on a local computing environment equipped with an AMD Ryzen 9 8945HX processor (2.50 GHz), 16 GB of RAM, and an NVIDIA GeForce RTX 5060 Laptop GPU running a 64-bit Windows 11 operating system. Under this hardware configuration, the framework required a total execution time of approximately 26.5 hours, yielding a weighted average processing time of 1.15 seconds per individual sequence.

The computational profile of NeuroCDS reflects its underlying neural-symbolic architecture. Compared to traditional heuristic tools or simple hexamer-based statistical models, the increased processing time is primarily attributed to the high-dimensional feature extraction within the dual-branch network. Specifically, the TCN module requires continuous evaluation of long-range codon usage frequencies across overlapping temporal windows, while the structurally constrained Viterbi algorithm involves maintaining a tripartite probabilistic state space across the entire sequence length.

Despite the additional neural overhead, the structurally constrained Viterbi algorithm operates with a strictly linear time and space complexity of *O*(*N*), where *N* is the sequence length. Because the state space size and evaluated reading frames are inherently small constants, and the state transitions are heavily restricted by unidirectional biological grammar, the decoding process completely avoids quadratic or exponential branching. This linear scaling ensures that NeuroCDS remains computationally feasible for large-scale *de novo* transcriptome projects. Furthermore, since the neural emission probability estimators are independent of the sequence parsing step, the framework can be significantly accelerated using modern GPU-based parallelization or distributed computing clusters. For most real-world applications, the trade-off between the marginal increase in computation time and the substantial gains in structural annotation robustness—especially for fragmented assemblies—represents a favorable balance for high-quality functional genomics.

## Discussion

The exceptional performance and robustness of NeuroCDS across diverse eukaryotic clades stem from its rational architectural design, which inherently mirrors the biological mechanisms of translation. Traditional tools often struggle because they rely on shallow statistical features or heuristic rules that fail under complex transcriptomic conditions. In contrast, our framework leverages a dual-branch deep learning strategy to independently capture distinct sequence signatures, thereby achieving a highly accurate initial probability estimation. Similar to the successful application of deep convolutional networks in specialized tools like TISRover [16], the CNN module in NeuroCDS serves as an effective local sensor, automatically extracting highly non-linear spatial motifs (such as the Kozak consensus) to precisely anchor Translation Initiation Sites. Concurrently, the TCN module evaluates the continuous regional coding potential. By utilizing an overlapping sliding-window strategy to extract a structured codon matrix, the TCN natively models synonymous codon usage biases. Crucially, as demonstrated in recent advancements like NeuroTIS+ [17], incorporating codon label consistency constraints within this temporal framework effectively suppresses intra-codon positional noise. This synergy between localized motif detection and smoothed regional coding evaluation provides an exceptionally high-precision initial probability landscape for the downstream decoding phase.

A primary algorithmic advantage of this neural formulation is its inherent resilience to sequence fragmentation, a pervasive bottleneck in *de novo* transcriptomics. Because the TCN relies on a sliding-window evaluation of the codon matrix, it extracts localized codon usage biases that are intrinsically distributed throughout the entire coding body. Consequently, the network does not depend on the global structural integrity of the transcript to accumulate continuous coding signals. When the framework encounters heavily truncated contigs—such as those completely lacking canonical start or stop codons—it can still stably delineate the active coding region based purely on the internal codon usage signatures captured within these overlapping windows. This “content-driven” resilience fundamentally explains why NeuroCDS successfully recovers functional information from information-poor partial fragments (e.g., NoStart/NoStop datasets), an area where traditional Markov models, which heavily rely on rigid transition paths anchored by defined global start and stop states, frequently falter.

Ultimately, the superiority of NeuroCDS is cemented by the integration of the Structurally Constrained Viterbi decoding algorithm, which transforms the isolated, probabilistic outputs of the deep neural networks into a biologically valid structural consensus. Purely neural models, despite their powerful pattern recognition capabilities, predominantly treat regional evaluation as isolated, point-wise prediction tasks. They inherently lack the capacity to enforce strict, sequence-level biological rules, occasionally yielding invalid outputs such as overlapping reading frames or unhandled internal stop codons. The Segmental Viterbi algorithm effectively resolves this critical limitation by mathematically fusing the heterogeneous deep features—the discrete spatial signals from the CNN and the continuous coding probabilities from the TCN—within a rigorous dynamic programming space. More importantly, it natively integrates fundamental biological structural constraints: the strict preservation of the triplet mRNA reading frame structure and the absolute avoidance of in-frame translation termination signals. By calculating the globally optimal path that satisfies both the neural emission probabilities and these physical grammars, NeuroCDS effectively bridges the representational power of modern deep learning with the mathematical interpretability of classical sequence parsing. This ensures that the final annotations are not only statistically highly accurate but also structurally flawless and immediately applicable for downstream functional genomics.

## Conclusion

In this study, we presented NeuroCDS, a reliable computational framework that effectively bridges the gap between the high-dimensional representation capabilities of deep learning and the structural rigor of classical dynamic programming. By mathematically coupling localized sequence features extracted via CNNs with global temporal coding signatures captured by TCNs, NeuroCDS provides an effective approach to mitigate the structural validity challenges often encountered in purely neural genomic models. Unlike previous end-to-end black-box architectures, our framework employs a Structurally Constrained Viterbi Decoding algorithm to enforce strict biological grammars—e.g., ensuring reading frame preservation—thereby calculating the globally optimal transcript structure in a joint inference space.

Comprehensive benchmarking demonstrates that NeuroCDS provides a highly adaptable solution for CDS annotation. It achieves high accuracy on intact eukaryotic transcripts and exhibits favorable robustness against the extreme fragmentation and strand ambiguity characteristic of *de novo* assemblies. Furthermore, the integration of a dynamic length normalization mechanism effectively neutralizes the deceptive pseudo-signals embedded within complex non-coding landscapes, such as 3’UTRs and ncRNAs. As the volume of transcriptome data continues to expand, particularly for non-model organisms, the interpretability and mathematical advantages of NeuroCDS offer a reliable foundation for downstream functional genomics, proteomics, and evolutionary biology.

## Key Points

- **Synergistic Dual-Branch Architecture:** NeuroCDS integrates Convolutional Neural Networks (CNN) for precise local motif extraction (e.g., translation initiation sites) and Temporal Convolutional Networks (TCN) for evaluating continuous regional coding potential based on synonymous codon usage biases.
- **Structurally Constrained Joint Inference:** Formulating CDS annotation as a holistic sequence parsing problem, the framework employs a Viterbi algorithm to systematically fuse heterogeneous deep features while strictly enforcing biological grammars, such as reading frame preservation and in-frame stop codon avoidance.
- **Robustness to Transcriptome Artifacts:** Driven by a sliding-window evaluation and dynamic length normalization, NeuroCDS exhibits exceptional resilience on heavily truncated contigs (e.g., NoStart/NoStop fragments), effectively recovering functional annotations even when canonical boundary signals are completely absent.
- **Stringent Specificity and Broad Generalization:** Extensive evaluations on complex Ribo-seq validated datasets and highly deceptive non-coding elements (e.g., ncRNAs and 3’ UTRs) confirm that the model delivers highly reliable, false-positive-resistant predictions across diverse eukaryotic clades.
- **Transparent and Highly Efficient Decoding:** Benefiting from unidirectional biological constraints, the dynamic programming engine operates with an optimized time complexity of *O*(*N*). This ensures strictly linear scaling for transcriptome-wide analysis while providing “glass-box” interpretability for visualizing the structural decision logic.

## Acknowledgments

The authors thank the anonymous reviewers for their valuable suggestions.

